# Gnotobiotic zebrafish microbiota display inter-individual variability affecting host physiology

**DOI:** 10.1101/2023.02.01.526612

**Authors:** Emmanuel E. Adade, Rebecca J. Stevick, David Pérez-Pascual, Jean-Marc Ghigo, Alex M. Valm

## Abstract

Gnotobiotic animal models reconventionalized under controlled laboratory conditions with multi-species bacterial communities are commonly used to study host-microbiota interactions under presumably more reproducible conditions than conventional animals. The usefulness of these models is however limited by inter-animal variability in bacterial colonization and our general lack of understanding of the inter-individual fluctuation and spatio-temporal dynamics of microbiota assemblies at the micron to millimeter scale. Here, we show underreported variability in gnotobiotic models by analyzing differences in gut colonization efficiency, bacterial composition, and host intestinal mucus production between conventional and gnotobiotic zebrafish larvae re-conventionalized with a mix of 9 bacteria isolated from conventional microbiota. Despite similar bacterial community composition, we observed high variability in the spatial distribution of bacteria along the intestinal tract in the reconventionalized model. We also observed that, whereas bacteria abundance and intestinal mucus per fish were not correlated, reconventionalized fish had lower intestinal mucus compared to conventional animals, indicating that the stimulation of mucus production depends on the microbiota composition. Our findings, therefore, suggest that variable colonization phenotypes affect host physiology and impact the reproducibility of experimental outcomes in studies that use gnotobiotic animals. This work provides insights into the heterogeneity of gnotobiotic models and the need to accurately assess re-conventionalization for reproducibility in host-microbiota studies.

## Introduction

Host-associated microbiota play essential roles in host physiology and health (1). Numerous studies have explored the perturbation of this microbiota in order to understand its contribution to host function, particularly for disease resistance, antibiotic effects, gut health, environmental factors, and many others (2–5). However, the complexity and heterogeneity of conventional host-associated microbiota, containing unculturable microorganisms and unknown functions, in humans and animal models limits mechanistic understanding.

In contrast to conventional animals with relatively complex microbiota (6), axenic (germ-free) and gnotobiotic animal models are powerful tools to study microbial contributions to host functions. These models allow for direct manipulation of bacterial genetics, composition, and exposure to determine the key drivers of microbiota-associated phenotypes under presumably more reproducible conditions. However, despite the capacity to control many aspects of gnotobiotic models, the inherent heterogeneity in biological systems could still contribute to phenotypic variability. In the context of host-associated microbiota, this could be due to pooled tissue samples, averaged replication, pseudo-replication within treatment conditions, and/or the use of simplified core microbiota (7,8). For example, previous studies of gnotobiotic mice have explored inter-facility microbiota variability (9) or colonization over time (10), but the samples were pooled prior to analysis and only presence/absence of taxa per group was considered.

Similarly, the study of host-pathogen-microbiota interactions in gnotobiotic zebrafish, an emerging model recapitulating many salient features of the mammalian gastrointestinal tract (11,12), is also subjected to variability. We indeed previously showed that reconventionalization of axenic zebrafish with bacterial strains isolated from the conventional fish provides variable protection against the fish pathogen *Flavobacterium columnare* (13). This suggests that gnotobiotic zebrafish reproducibly reconventionalized with a controlled bacterial consortium is prone to underreported inter-individual variability that may be the origin of phenotypic variation.

Here we explored this issue by investigating intra and inter-individual microbial composition, colonization biogeography and host mucus production as an indicator of host physiology in conventional and gnotobiotic zebrafish models. We generated gnotobiotic animals using a previously-studied consortium of 9 bacterial strains isolated from the conventional fish microbiota (13). We showed that larval zebrafish gut bacterial load fluctuates and that bacterial colonization, composition and biogeography varies between conventional and gnotobiotic fish models, which impacts zebrafish mucus production and tissue architecture. While providing insights on baseline variability of gnotobiotic models harboring bacterial communities of reduced complexity under control conditions, our study also illustrates the necessity to characterize this variability before investigating the effects of antibiotics, disease, or other perturbations on host-associated microbiota.

## Results

### Bacterial colonization of zebrafish larvae fluctuates and is localized to the gut

In order to investigate inherent variability between conventional zebrafish and zebrafish reconventionalized with the previously-studied consortium of 9 bacterial strains (Mix9) isolated from the conventional fish microbiota (13), we measured the bacterial biovolume in zebrafish tagged with the universal bacterial probe Eub338 (14) (**Fig 1A-D**). We observed significantly higher bacterial colonization in the conventional larvae than in the Mix9 fish, with median bacteria biovolume of 2.7 × 10^5^ μm^3^ and 2.6 × 10^4^ μm^3^ respectively (p-value = 0.035) (**Fig 1E**). These bacterial biomasses were primarily localized to the intestinal region of the larval zebrafish, with a smaller number of cells colonizing the skin (**Fig 1A-D**). Consistent with sterility tests performed throughout the study, no bacteria were detected in axenic (germ-free) zebrafish (**Fig 1E**). We then analyzed a larger sample of 24 reconventionalized Mix9 larvae, and we observed a high variability in bacterial biovolume among experimental animals as measured with confocal imaging (**Fig S1**). CFU counting corresponding to entire homogenized zebrafish plated on agar growth medium confirmed the variability observed in the FISH biovolume measurements (**Fig 1F**).

**Figure 1.**
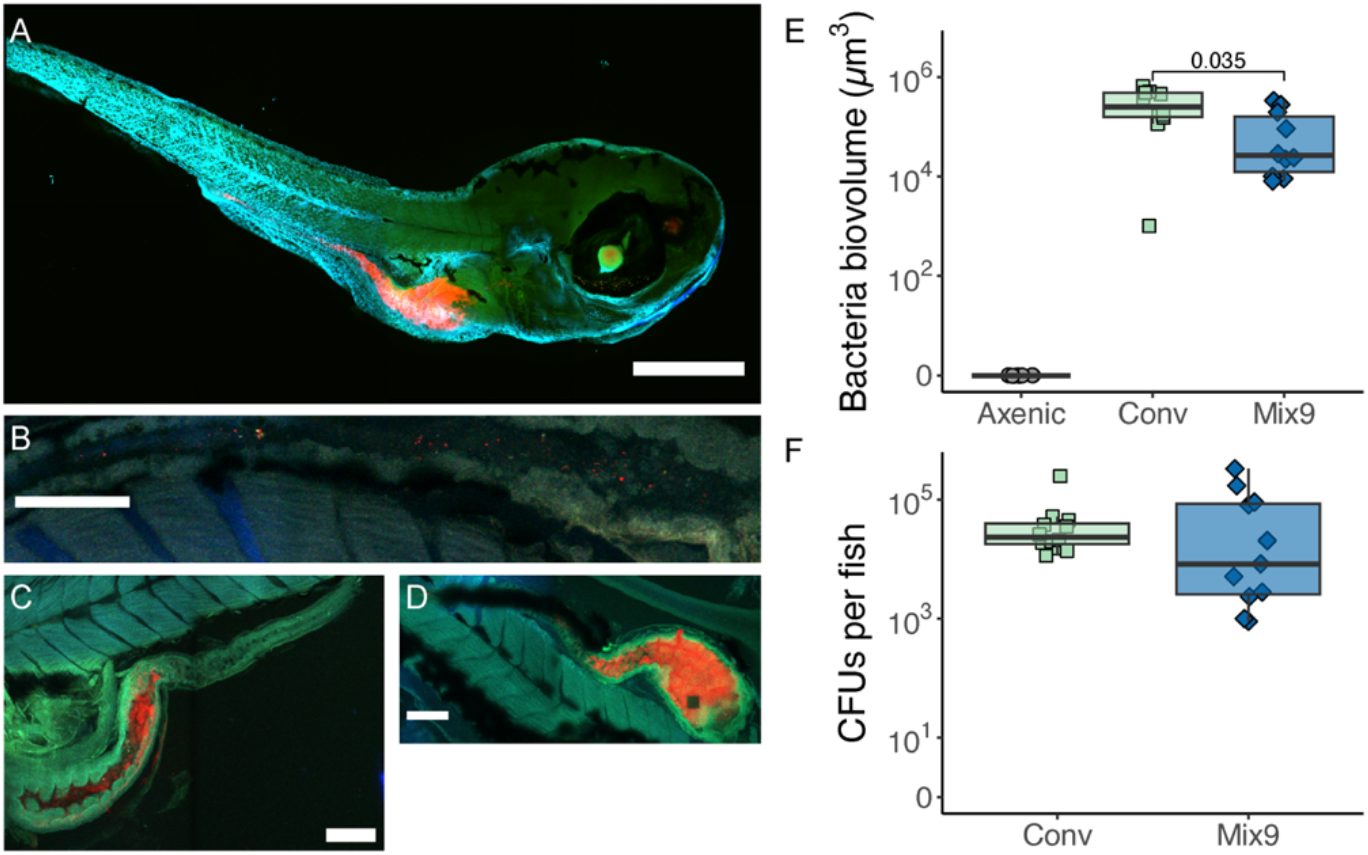
Bacterial load was localized in the larval zebrafish gut and showed colonization variability. (**A**) Detection of bacteria by Fluorescent in situ Hybridization (FISH) on fixed zebrafish larvae exposed to Mix9 at 7 days post fertilization (dpf) showed the bacteria localized primarily in the gut. Representative images of (**B**) zebrafish gut with low bacterial load, (**C**) zebrafish gut with medium bacterial load, and (**D**) zebrafish gut with high bacterial load. (**A-D**) EUB 338 conjugated with Alexa Fluor 594 (Red)– All bacteria, DAPI (Blue) – (host nuclei), Zebrafish autofluorescence in (Green). Scale bars (A) = 200 μm, (B, D) = 100 μm and (C) = 40 μm. (**E**) Bacterial biovolume as a measure of abundance with FISH. Unpaired T-test (alpha = 0.05) was used to measure significance in variation between conventional and Mix9 (n = 10). (**F**) Quantification of bacterial load in whole larval zebrafish (Conventional and Mix 9) via culturable CFUs. Bacterial load quantification via CFU counts of homogenized Mix9 fish on LB media at 7 dpf (n = 10). Note log scale on y-axis for **(E)** and **(F)**.

### Bacterial colonization biogeography varies between conventional and gnotobiotic models

In addition to the bacterial load, we also examined the bacterial biogeography per fish and observed that there is variability in the bacterial localization along the length of the intestinal tract (**Fig 2A**) in both conventional and gnotobiotic Mix9 zebrafish larval models (**Fig 2B**). Conventional larvae were characterized by a consistently high bacterial load in the bulb and less in the proximal intestine and on average lower but heterogenous bacterial load in the distal intestine (**Fig 2D**). However, Mix9 larvae had a highly heterogeneous bacterial load in the bulb among animals ranging from no bacteria to approximately 1 × 10^5^ μm^3^ of bacteria, and more consistent biovolumes in the proximal and distal intestines (**Fig 2E**).

**Figure 2.**
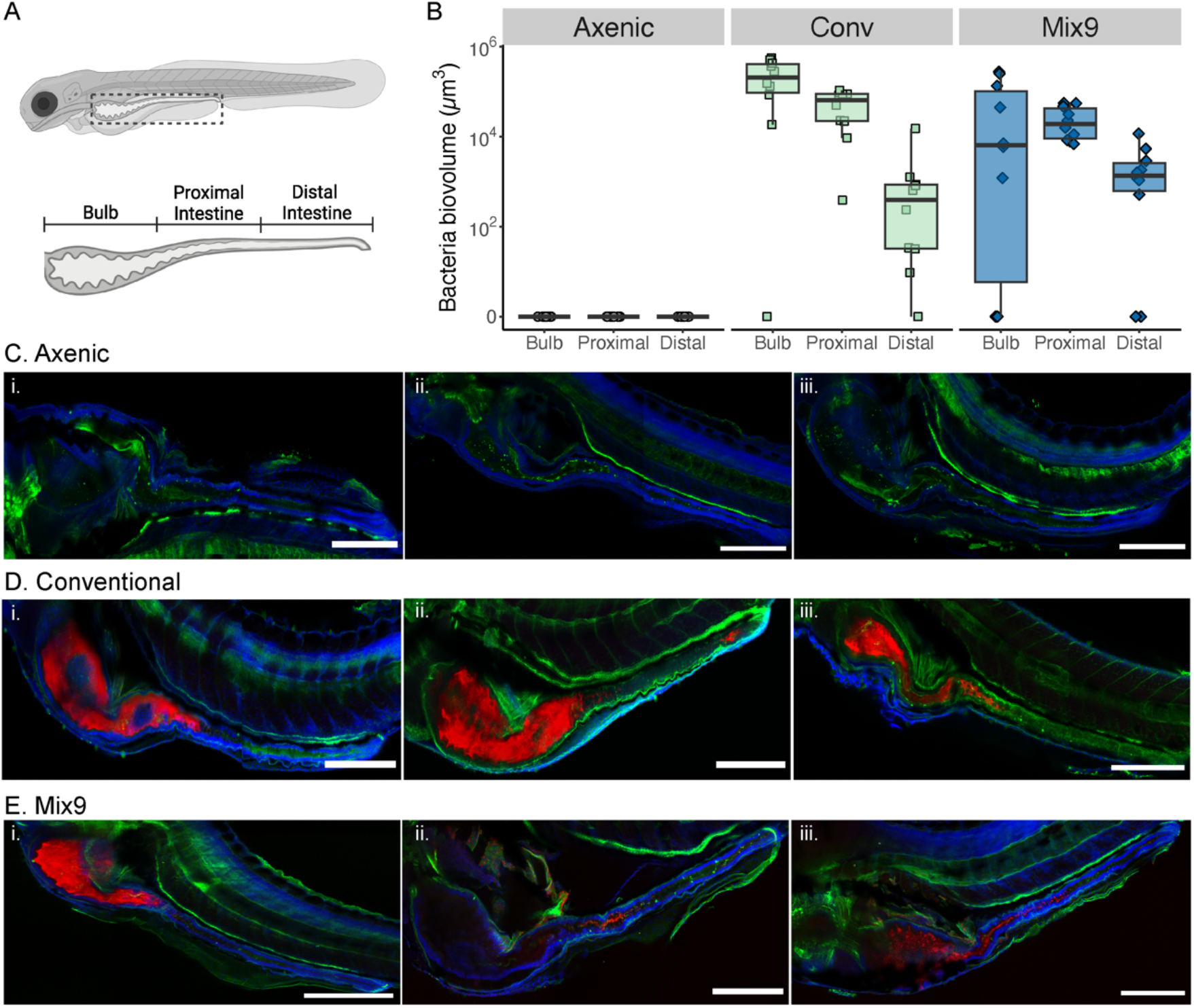
Bacterial biogeography and abundance along the gastrointestinal tracts of the larval zebrafish. (**A**) A schematic of the larval stage zebrafish with the gut highlighted. [Inset] The schematic gut of the larval zebrafish with the segmentation classifying the three sections of the gut analyzed here. (**B**) Analysis of the spatial distribution of bacteria in the gut for the various conditions (n=10). (**C – E**) Confocal images of representative larvae in each condition shown. **i, ii** and **iii** show three representative images for each condition. Universal bacteria probe EUB 338 conjugated with Alexa Fluor 594 (Red)– All bacteria, DAPI (Blue) – host nuclei, WGA conjugated with Oregon Green 488 (Green) – Mucus in the fish. All scale bars = 200 μm.

### Microbiota in conventional zebrafish larvae is more diverse than in gnotobiotic larvae

We further evaluated the differences in bacterial composition between conventional and gnotobiotic zebrafish using 16S rRNA gene amplicon sequencing (**Fig S2**). Despite the reduced community administrated to the gnotobiotic Mix9 zebrafish, their composition at the phylum level is not significantly different from the conventional fish (**Fig 3A**). Both the conventional and Mix9 fish bacterial composition is dominated by > 90 % Proteobacteria, with the remainder in the Bacteroidota and Actinobacteria phyla. At the Amplicon Sequence Variant (ASV) level, the conventional fish microbiome is mainly comprised of a single *Aeromonas* sp. ASV 02, with over 50 % relative abundance in each sample (**Fig 3B**). There was a relative decrease in *Aeromonas* sp. ASV 02 and an increase in ASV 01 unclassified Gammaproteobacteria (likely *Pseudomonas sp*.) in the Mix9 condition, compared to conventional fish (**Fig 3B**). Despite addition of 9 bacterial strains in equal proportion to the axenic fish model at 4 dpf, only two ASVs were identified in all fish at 7 dpf: 02 *Aeromonas* sp. and 01 Gammaproteobacteria *Pseudomonas* sp. These two ASVs may comprise multiple strains of the Mix9 that are indistinguishable at the 16S rRNA level.

**Figure 3.**
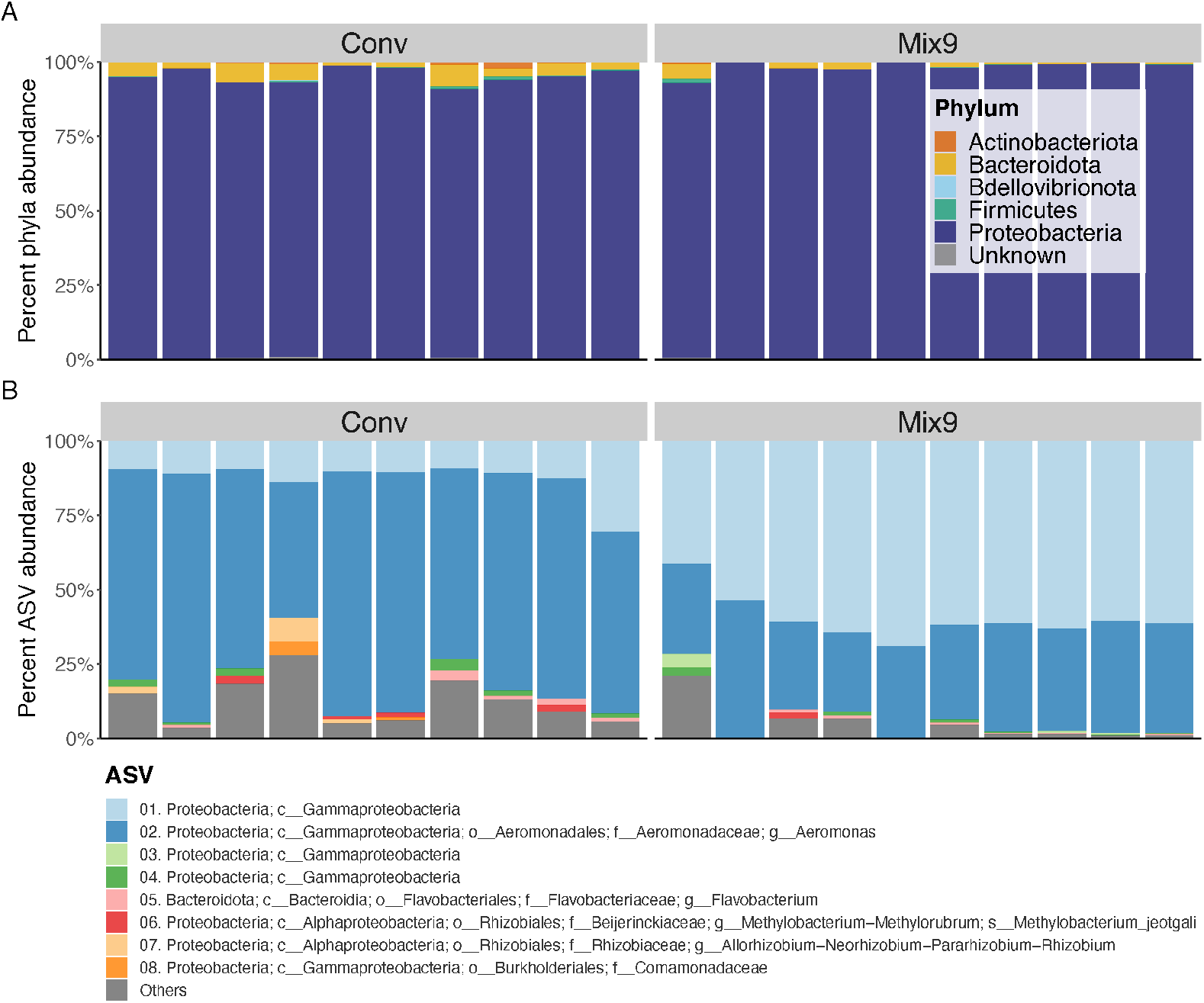
Bacterial composition of conventional and Mix9 fish at the (A) Phylum and (B) ASV level. Each stacked bar represents an individual larval fish sampled at 7 dpf with n = 10 individual fish per condition. **(A)** Bar plots of percent phylum abundance per sample. **(B)** Bar plots of percent Amplicon Sequence Variant (ASV) abundance per sample. The top 8 most abundant ASVs are shown with the others grouped into “Others.”

Conventional and Mix9 fish shared 32 ASVs, while conventional fish had 54 unique ASVs (55% of total ASVs) (**Fig 4A**). Despite many shared taxa, the conventional fish microbiota had a significantly higher alpha-diversity than the Mix9 fish microbiota as measured by Chao and Shannon indices, but not Simpsons index (**Fig 4B**). Simpson’s index has decreased sensitivity to less abundant taxa, indicating that the differences between the two models is in the rare taxa and that the most abundant taxa are comparable. This is further demonstrated with no significant difference in beta-diversity or beta-dispersion between the conventional and Mix9 microbiota (**Fig 4C-D**). Overall, the conventional fish have a higher richness and increased number of less abundant taxa than the simplified microbiota introduced in the Mix9 gnotobiotic fish.

**Figure 4.**
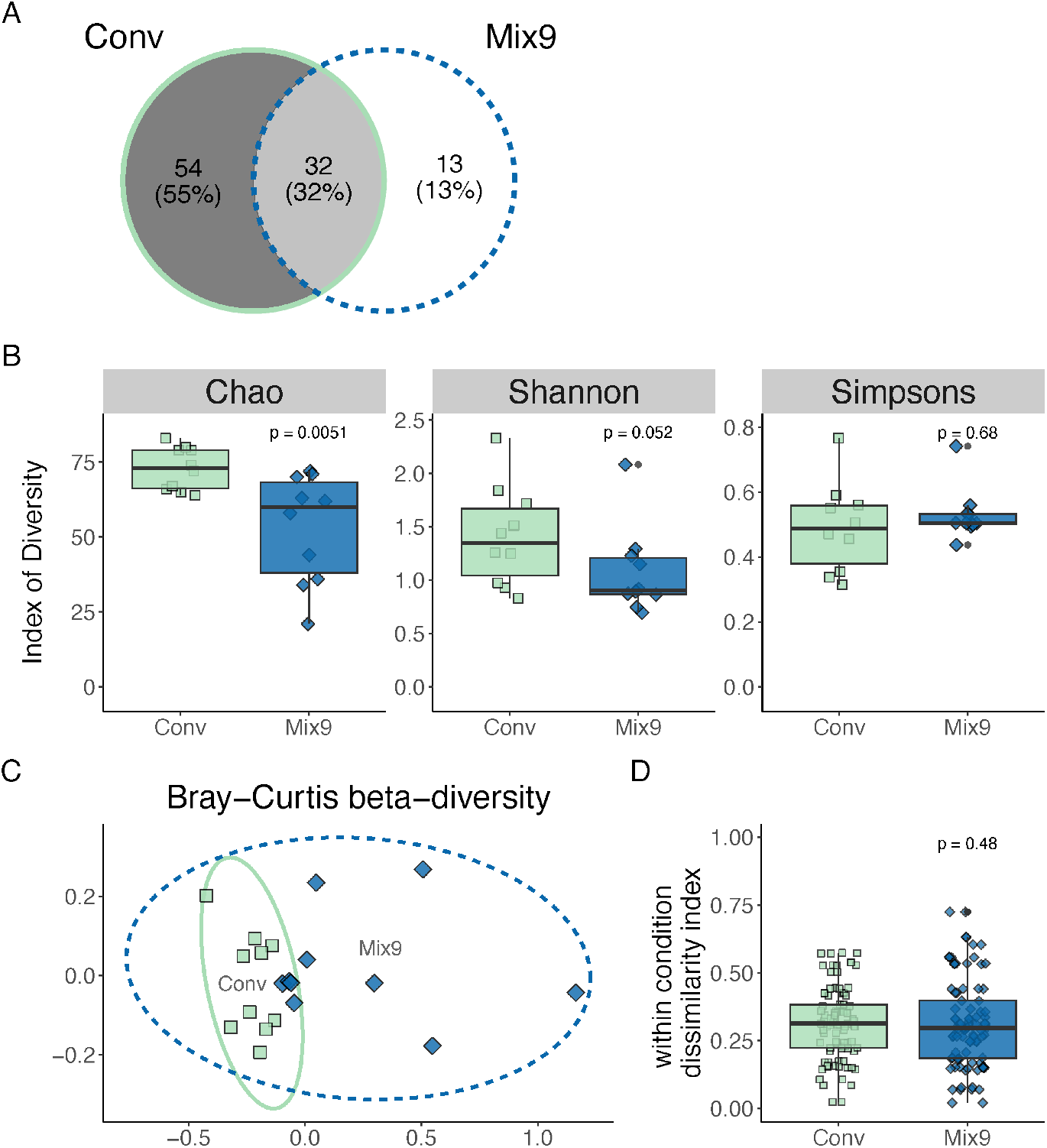
Alpha- and beta-diversity of individual conventional and Mix9 fish bacterial communities measured using 16S rRNA amplicons. **(A)** Venn diagram of ASVs shared between conventional (Conv) and Mix9 conditions. All ASVs with less than 5 reads were removed. **(B)** Alpha-diversity calculated using Chao richness, Shannon, and Simpson’s indices. **(C)** NMDS plot calculated using Bray-Curtis beta-diversity (k=2) of percent normalized ASVs for conventional and Mix9 samples. Ellipse lines show the 95 % confidence interval (standard deviation). Stress = 0.055. **(D)** Beta-dispersion or within-condition dissimilarity index calculated using Bray-Curtis beta-diversity. **For all panels:** n = 10, Wilcoxon test for Mix9 compared to the Conventional (Conv) condition.

### Changes in zebrafish mucus production and tissue architecture depend on microbiota composition and colonization

Mucus, composed mostly of the highly O-glycosylated mucins, coats the epithelial surfaces of the gastrointestinal tract and has important functions, including the prevention of colonization by foreign microbes (15–20). The interactions among resident gut microbes and between microbiota and host has been shown to influence mucus production and shape microbiota spatial distribution on a macroscale (21). To better understand mucus biogeography and its potential variability, we monitored the presence of gut mucus in the conventional, axenic and Mix9 zebrafish models with fluorescent wheat germ agglutinin (WGA) staining (**Fig 5A-D**).

**Figure 5.**
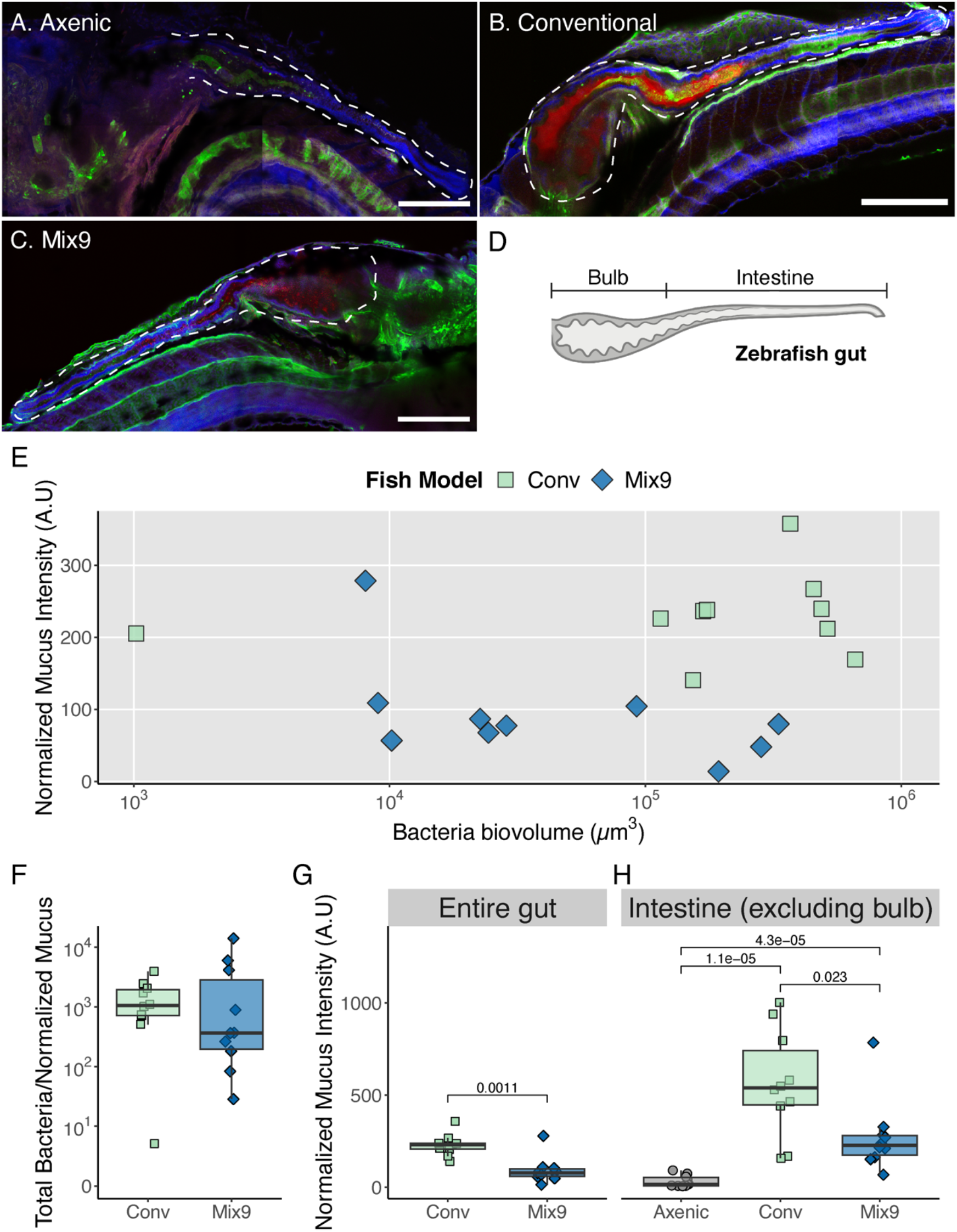
Zebrafish intestinal mucus production in multi-member gnotobiotic model is different than conventional fish. **(A – C)** Confocal images of the gut region of the zebrafish for each condition. Universal bacteria probe EUB 338 conjugated with Alexa Fluor 594 (Red) – All bacteria, DAPI (Blue) – (host nuclei), WGA conjugated with Oregon Green 488 (Green) – Mucus in the fish, (Magenta) – Zebrafish autofluorescence. All scale bars = 200 μm. **(D)** The schematic larval zebrafish gut with the segmentation classifying the two sections of the gut analyzed here. **(E)** Per fish correlation of total bacteria load and normalized mucus intensities. **(F)** The relative measurement of mucus to bacteria biovolume per fish shown on a log scale. **(G-H)** Normalized mucus intensity in the entire gut and intestine excluding bulb. For entire gut, T-tests were used to estimate the statistical significance (alpha = 0.05), p-values < 0.05 were significant. For intestine only, One-Way ANOVA and Tukey post hoc test was used to statistical measure the variation among the various conditions, p-values < 0.05 were significant. (n = 10).

Intriguingly, we observed little correlation between total bacterial load and mucus abundance, per animal, in the conventional and Mix9 models (**Fig 5E-F**). This suggests that induction of mucus production by the gut microbiome may be more dependent upon the presence and distribution of specific organisms, rather than the total bacterial biomass. Comparison of mucus levels in all 3 models focused on the intestine only, because the bulb tissue architecture was degenerated in the axenic control, and therefore difficult to quantify with imaging. We observed that mucus abundance was highest in conventional fish, intermediate in Mix9 and lowest in axenic (**Fig 5G-H**). Histological staining with Alcian blue and Periodic-Acid Schiff (PAS) stains performed at 7 dpf (72 h post-reconventionalization) on axenic, conventional, and Mix9 gnotobiotic larvae did not show any major qualitative difference in their intestinal organization (**Fig S3**).

Together these data suggest that bacterial colonization and composition affect mucus production without any impact on the tissue architecture, leading to changes in host physiology, with the highest variability in the Mix9 gnotobiotic condition.

## Discussion

In this study, we explored the inter-individual relationship between host bacterial colonization, composition, and mucus production as an indicator of host physiology. Whereas the conventional zebrafish showed higher taxonomic diversity than the Mix9 gnotobiotic zebrafish, their microbiota beta-diversity was not significantly different. Nevertheless, our analysis revealed a remarkable heterogeneity in the distribution of bacteria along the length of the gut in the gnotobiotic model. Heterogeneity in the biogeography and abundance of bacteria along the longitudinal (mouth to anus) and transverse (epithelium to lumen) axes of the gut is known to be driven by numerous host, microbial and environmental factors. This includes peristaltic activity, microbial motility, mucus architecture, chemical and nutrient gradients, host immunity and both synergistic and antagonistic bacterial interactions (22,16,23–26). Intriguingly, we observed bacteria as individual cells, small aggregates, and large clusters in the gut in our Mix9 gnotobiotic model.

As previously reported, mucus was generally more abundant in the bulb and proximal region of the zebrafish gut (20), but its overall abundance was highest in conventional larvae, intermediate in Mix 9 and lowest in axenic larvae. Similar observations were made in axenic (germ-free) mice, in which mucus was shown to differ in the small and large intestine, and to depend on their bacterial composition (27–29). These results are therefore indicative of a dynamic interrelationship between microbiota and host mucus production that is not only due to the presence or absence of a microbiota but results more from subtle spatial relationships between specific microbial taxa and the host. We hypothesize that there may be different combinations of bacteria species locally present in each fish gut in a spatially dependent manner. Further analysis of gut biogeography and architecture with higher phylogenetic resolution may provide support for this hypothesis.

The reduced complexity of our gnotobiotic zebrafish model enabled us to observe that the Mix9 bacterial consortium restored bulb tissue architecture, including mucus production, that otherwise deteriorated in axenic animals. Interestingly, our histological analysis showed that the gut tissue of axenic and Mix9 fish appeared morphologically no different to conventional ones, consistent with previously reported phenotypic analysis of gnotobiotic zebrafish (30,31). These studies also reported a significant decrease in the number of Goblet cells in axenic zebrafish larvae compared to conventional ones (30). These discrepancies could originate from the different feeding protocols and the use of sterile powder food versus sterile live *T. thermophila* in our study (31) or even not fed at all (30).

Despite the use of autochthonous bacterial strains isolated from conventional zebrafish larvae, we observed a high variability in bacterial load as well as in microbiota and spatial structure in gnotobiotic Mix9 larvae compared to conventional fish. Whereas this was previously observed in zebrafish larvae mono-reconventionalized with *Pseudomonas* sp. or *Aeromonas* sp., strains (30), our data identified an important limitation as gnotobiotic animals reconventionalized with low complexity microbiomes exhibit heterogeneous colonization efficiencies and mucus phenotypes not observed in conventional animals, which may have profound implications for reproducibility.

Various phenomena could affect the assembly of the microbiota in our gnotobiotic model, including differences in bacterial migration into the fish, survival to the gut environment, bacterial interactions, intestinal expulsion, or predation by live *Tetrahymena* used as zebrafish food source (32). In addition, the administration of bacteria in fish water and their acquisition by natural routes may contribute to the observed variability in colonization (12). The use of direct and more controlled inoculation methods including microinjection or gavage could reduce colonization heterogeneity.

Gnotobiotic animal models are invaluable tools to study host-associated microbiota function. However, our study demonstrates that special attention should be given to key parameters such as bacterial load and composition before assuming control and reproducibility of the results obtained with these gnotobiotic models (33). These criteria should be taken into consideration as they are likely to impact colonization efficiency as well as effects on the host and future studies of the factors stabilizing host-microbiota interactions will contribute to validate and increase the usefulness of gnotobiotic approaches.

## Methods

### General zebrafish husbandry

Wild-type AB/AB zebrafish (*Danio rerio*) fertilized eggs at 0 days post fertilization (dpf) were obtained from the Zorgl’hub platform at Institut Pasteur. All procedures were performed at 28°C under a laminar microbiological hood with single-use plastic ware according to European Union guidelines for handling of laboratory animals and were approved by the relevant institutional Animal Health and Care Committees. Eggs were kept in 25 cm^3^ vented flasks (Corning 430639) with 20 mL of autoclaved mineral water (Volvic) until 4 dpf (30 – 33 eggs/flask), transferred to new flasks after hatching at 4 dpf (10 – 15 fish/flask), then transferred to individual wells of a 24-well plate (TPP 92024) at 6 dpf. At the end of the experiment, zebrafish were euthanized with tricaine (MS-222, Sigma-Aldrich E10521) at 0.3 mg/mL. Fish were fed with sterile *T. thermophila* every 48 hours starting at 4 dpf as previously described (34).

### Zebrafish sterilization and reconventionalization

The zebrafish embryos were sterilized as previously described at 1 dpf and then maintained as described above (13). Sterility was confirmed at 3 dpf by spotting 50 μL of water from each flask on LB, TYES and YPD agar plates and incubated at 28 °C under aerobic conditions for at least 3 days. Contaminated flasks were immediately removed from the experiment and not included in the results. Axenic zebrafish larvae were reconventionalized at 4 dpf, as follows. Overnight cultures of a single bacterial colony in 5 mL of liquid media were washed twice with sterile mineral water (Volvic) and normalized to OD-0.1 in water. For Mix9 reconventionalization, 200 μL of each strain was added per flask at a final concentration of 5 × 10^5^ CFU/mL per strain. Water samples were plated in serial dilutions to confirm final bacterial concentration and sterility. Reconventionalization was performed for 48 hours until fish were transferred to sterile water in 24-well plates at 6 dpf.

### Bacterial strains and growth conditions

Bacterial strains are listed in Table 1. All strains were grown in Miller’s Lysogeny Broth (LB) (Corning) and incubated at 28°C with rotation. Cultures on solid media were on LB with 1.5 % agar. Bacteria were always streaked from glycerol stocks stored at -80°C on LB--agar before inoculation with a single colony in liquid cultures. All media and chemicals were purchased from Sigma-Aldrich.

**Table 1.** Bacterial strains used in this study. All strains were isolated from conventional zebrafish larvae (13).

| Code | Strain |
| --- | --- |
| UGB3608 | *Aeromonas veronii* 1 |
| UGB3612 | *Pseudomonas mossellii* |
| UGB3616 | *Stenotrophomas maltophilia* |
| UGB3607 | *Aeromonas caviae* |
| UGB3614 | *Pseudomonas peli* |
| UGB3615 | *Pseudomonas sediminis* |
| UGB3611 | *Phyllobacterium myrsinacearum* |
| UGB3609 | *Aeromonas veronii* 2 |
| UGB3613 | *Pseudomonas nitroreducens* |

### Quantification of zebrafish bacterial load via CFU counts

Zebrafish were sampled at 7 dpf and washed twice by 2 transfers to clean, sterile water in petri dishes to remove loosely attached bacteria. The larvae were then added in 500 μL of sterile water to 2 mL tubes containing 1.4 mm ceramic beads (Fischer Scientific 15555799) and homogenized for 2 × 45 seconds at 6000 rpm using a 24 Touch Homogenizer (Bertin Instruments). These homogenization conditions are sufficient to lyse zebrafish tissue, but not harmful to the bacteria. The lysate was then diluted from 10- 100-fold and 3 × 100 μL from each dilution was spread on LB agar using sterile glass beads. After 2 days of incubation at 28°C, CFUs were counted per fish and calculated by 500 μL lysate / 100 μL plated * dilution factor * (average of replicate CFUs).

### Zebrafish sampling and DNA extraction

At 7 dpf, 10 zebrafish of each conventional and Mix9 conditions were sampled and washed twice by 2 transfers to clean, sterile water in petri dishes. Each fish was then added to a sterile 2-mL microcentrifuge tube and euthanized with tricaine at 0.3 mg/mL. All liquid was removed from the tissue and the samples were immediately frozen at -80°C until DNA extraction. DNA extraction was performed from single larval zebrafish using the DNeasy Blood & Tissue kit (Qiagen 69504) with modifications as follows. Tissue samples were thawed at room temperature, then 380 μL Buffer ATL and 20 μL proteinase K were added directly to each individual larva in a 2 mL tube. Samples were vortexed, then incubated overnight (15-18 hours) at 56°C and 300 rpm until fully lysed. After lysis, 4 μL of RNAse A solution was added and the samples were incubated for 5 minutes at room temperature to remove residual RNA. Next, 400 μL Buffer AL and 400 μL 100 % ethanol were added and mixed by vortexing before loading the lysate onto the DNeasy mini spin column in 2 × 600 μL loads. DNA purification and cleanup proceeded according to the manufacturer’s recommendations with a final elution volume of 50 μL in Buffer AE. Purified DNA was quantified using the Qubit HS DNA fluorometer kit (ThermoFisher Q32851) and purity was assessed with the Nanodrop spectrophotometer (ThermoFisher). Negative controls for the extraction kit were prepared alongside zebrafish samples, but with no tissue input.

### Zebrafish 16S rRNA amplicon sequencing and analysis

16S rRNA gene amplicons of the V6 region for the 10 conventional zebrafish samples, 10 Mix9 zebrafish samples, 2 mock community samples (Zymo Research DNA standard I D6305), 2 negative DNA extraction samples, and blank PCR control were prepared using 967F/1064R primers. A two-step PCR reaction using 200 ng of zebrafish DNA was performed in duplicate 50 μL reactions as previously described (35,36). Each first step reaction included 25 μL 2X Phusion Mastermix (Thermo Scientific F531S), 1.5 μL of 10 μM F/R primer mix, 13 - 20 μL template DNA (200 ng), and 3.5 – 10.5 μL nuclease-free water (up to 50 μL). PCR amplification (step 1) conditions were denaturing at 98°C for 3 min followed by 30 cycles of denaturation at 98°C for 10 s, primer annealing at 56°C for 30 s, and extension at 72°C for 20 s, then a final extension at 72°C for 20 s. Negative controls for the PCR reagents were prepared alongside zebrafish DNA samples, but with additional nuclease-free water input. PCR products were assessed for concentration (Qubit DNA HS reagents) and expected size using agarose gel electrophoresis. A second PCR step was performed to attach sequencing barcodes and adaptors according to Illumina protocols. The PCR products were analyzed with 250 bp paired-end sequencing to obtain overlapping reads on an Illumina MiSeq at the Institut Pasteur Biomics platform.

The resulting 16S rRNA gene amplicon sequences were demultiplexed and quality filtered using DADA2 (v1.6.0) implemented in QIIME2 (v2020.11.1) with additional parameters --p-trunc-len-r 80 --p-trunc-len-f 80 --p-trim-left-r 19 --p-trim-left-f 19 to determine amplicon sequence variants (ASVs) (37,38). All ASVs were summarized with the QIIME2 pipeline (v2020.11.1) and classified directly using the SILVA database (99 % similarity, release #134) (39,40). Processed ASV and associated taxonomy data was exported as a count matrix for analysis in R (v4.1.3). The positive and negative controls were checked to ensure sequencing quality and expected relative abundances. Non-bacterial and chloroplast sequences were then removed, and the data was normalized by percentage to the total ASVs. All ASVs with less than 0.1% abundance were removed from each sample for further dissimilarity metric analysis (41).

All descriptive and statistical analyses were performed in the R statistical computing environment with the *tidyverse* v1.3.1, *vegan* v2.5.7 and *phyloseq* v1.38.0 packages (42–44). Non-metric dimensional analysis (NMDS) was used to determine the influence of fish type on the ASV-level composition. The Bray-Curtis dissimilarity metric was calculated with k = 2 for max 50 iterations and 95 % confidence intervals (standard deviation) were plotted. Statistical testing of the beta-diversity was done using the PERMANOVA *adonis2* test implemented in *vegan* (method = “bray”, k = 2) (45,46). Within-condition variability was calculated using the command vegdist(method = “bray”, k = 2) and the matrix was simplified to include samples compared within each timepoint. Alpha-diversity metrics were calculated for each sample at the ASV level using the *vegan* package and analyzed using the non-parametric Kruskal–Wallis rank sum test in R. All processed sequencing files, bash scripts, QIIME2 artifacts, and Rmd scripts to reproduce the figures in the manuscript are available on Zenodo (47).

### Estimation of zebrafish bacterial load via FISH and mucus labelling

Whole larval zebrafish reconventionalized with Mix9 were preserved with Carnoy’s fixative 24 hours after infection (72 hours after reconventionalization) and bacteria were labelled using the Eub338 general probe by fluorescent in situ hybridization (FISH). The fish samples were imaged using confocal fluorescent microscopy to localize and quantify the bacterial load in each fish. Labeling of zebrafish was carried out in Eppendorf tubes. Carnoy’s fixed zebrafish samples were washed in PBS for 3 mins. For the bacteria labelling, to each tube, 200 μl of hybridization buffer [0.09 M NaCl, 0.02 M Tris pH 7.5, 0.01 % SDS, 20 % formamide] and 2 μl of EUB 338 FISH probe (GCTGCCTCCCGTAGGAGT) conjugated to Alexa Fluor 594 (Thermofisher) were added to the samples and incubated at 46 °C overnight (18-24 hrs). Fish were washed in 500 μl of wash 1 [0.09 M NaCl, 0.02 M Tris pH 7.5, 0.01 % SDS, 20 % formamide] for 30 mins at 48 °C followed by 1000 μl of wash 2 [0.09 M NaCl, 0.02 M Tris pH 7.5, 0.01 % SDS] for 30 mins at 48 °C and resuspended in 500 μl of resuspension buffer [0.02 M Tris, 0.01 % SDS] for 30 mins at RT in the dark. For mucus labelling, the resuspension buffer was replaced by 200 μl of 40 μg/ml Oregon Green labeled Wheat Germ Agglutinin (WGA) (Thermofisher) for 25 mins at RT in the dark. The labelling solution was removed, and samples were further washed twice in resuspension buffer for 5 minutes each. Samples were incubated in 200 μl of 0.55 μM DAPI for 10 mins at RT. Samples were washed twice with resuspension buffer. Samples were mounted on Ultrastick glass slides. Fish were oriented so larvae will be on its side. Specimens were mounted in Vectashield anti-fade reagent (Vector Laboratories).

### Zebrafish confocal image acquisition and pre-processing

A Zeiss LSM 710 confocal microscope (Carl Zeiss) was used to acquire spectral images. All images were acquired with Plan-Apochromat 20x 0.8 NA objective in lambda mode with 29.1 nm channel bandwidth resulting in 9 channels detected over the visible spectrum. Simultaneous excitation with 405 nm, 488 nm and 561 nm lasers was used. 3-D z-stack images were acquired as tile scans with 9 z-planes centered in the middle of the gut spanning 8.3 μm total in z dimension. Linear unmixing was performed on the spectrally acquired images after stitching the tiles together in ZEN software v 3.4. We extracted the reference spectra for DAPI, EUB 338 Alexa flour 594, WGA Oregon green 488 and autofluorescence from the labelled zebrafish and applied them for the linear unmixing.

### Image quantification and analyses

The unmixed images were further processed and analyzed in IMARIS v 9.6.1 (Bitplane) to quantify the bacteria biovolume and mucus intensity. For each larva, measurements were made separately in 3 gastrointestinal regions: the bulb, proximal and distal gut. The bulb region was defined using visual identification of the pylorus. The remainder of the GI tract, the intestine, was divided into two equal halves to define the proximal and distal gut in this analysis. For each of the three manually segmented regions 3D surfaces were generated for the labelled bacteria, using the default threshold algorithm in IMARIS v. 9.6.1 (Bitplane). After rendering surfaces, total volume measurements were calculated for bacteria.

To quantify mucus, the gut was manually segmented in each of the 9 z-planes for each larva using the columnar epithelium DAPI signal as a guide, because WGA labels both mucus as well as cell membrane glycoproteins present throughout zebrafish tissues. The WGA channel for the manually segmented region of interest in the gut lumen. Total intensity of WGA label in that area in each of the 9 z-planes was measured, then divided by the surface area of the manually segmented gut region of interest.

### Imaging statistical analyses

Students t-test was used to compare normalized mucus intensity between the conventional and Mix9 conditions. ANOVA (parametric) with Tukey post hoc test was used to compare variations in normalized mucus intensity in the intensity for the axenic, Mix9 and conventional conditions. For the comparison of the total bacteria biovolume. All analyses were done in R (v4.1.3).

### Histological analysis of zebrafish tissues

Histological sections were used to compare microscopical tissular organization between Conv, GF and mix9 zebrafish larvae. A total of 5 fish were sacrificed and fixed for 1 day in Carnoy’s fixative. Whole fixed animals were then dehydrated in methanol (2 × 30 minutes) then in ethanol 100 % (2 × 20 min). Final dehydration was performed by 100 % xylene solution 2 × 2 hours. Then, samples were embedded in paraffin wax solution (3 × 2 hours) and embedded in paraffin wax for polymerization. Sections (thickness 5μm) were cut with a microtome RM2245 (Leica Microsystems GmbH, Wien, Austria), and mounted on adhesive slides (Klinipath-KP-PRINTER ADHESIVES). Paraffin-embedded sections were deparaffinized and stained with Alcian Blue (AB) and Periodic-Acid Schiff (PAS) to observe both neutral and acidic mucins and Goblet cells quantification. All slides were scanned with the Panoramic Scan 150 (3D Histech) and analyzed with the CaseCenter 2.9 viewer (3D Histech). Goblet cells quantification was estimated by manual counting of total AB positive cells in blue per villi of the posterior gut.

## Acknowledgements

We are grateful to Bianca Audrain for assistance with obtaining the animal ethical authorization. Zebrafish embryos were obtained from Sebastian Bedu at the Zorgl’hub platform at Institut Pasteur. This work was supported by U.S. National Institutes of Health Grant R01DE030927 to AMV, the French Government’s Investissement d’Avenir program, Laboratoire d’Excellence “Integrative Biology of Emerging Infectious Diseases” (grant n°ANR-10-LABX-62-IBEID) to JMG, the Fondation pour la Recherche Médicale (grant DEQ20180339185) to JMG, and a grant from the Philippe Foundation to RJS. Sequencing was performed by G M. Haustant, L. Lemée, Biomics Platform, C2RT, Institut Pasteur, Paris, France, supported by France Génomique (ANR-10-INBS-09-09) and IBISA.

## Author Contributions

All authors contributed conception and design of the study. EA performed all microscopy and image analysis, with assistance from RJS and AMV. RJS and DPP performed the zebrafish experiments. RJS performed the 16S rRNA gene amplicon study and analyzed the data. JMG and AMV acquired funding for the study. All authors contributed to manuscript revision, read and approved the submitted version.

## Ethics Statement

All animal experiments described in the present study were conducted at the Institut Pasteur according to European Union guidelines for handling of laboratory animals (http://ec.europa.eu/environment/chemicals/lab_animals/home_en.htm) and authorized by the Institut Pasteur institutional Animal Health and Care Committees under permit #dap200024.

## Data Availability

The raw 16S rRNA gene amplicon sequences generated for this study can be found in the NCBI Sequencing Read Archive in <u>BioProject no. PRJNA928247</u>. All other raw data, processed sequencing files, and scripts to reproduce the figures in the manuscript are available in the Zenodo repository, https://doi.org/10.5281/zenodo.7573109 (47).

