## Supplementary Figures for "Gnotobiotic zebrafish microbiota display inter-individual variability affecting host physiology"

This PDF file includes:

Supplementary Figures S1 to S3

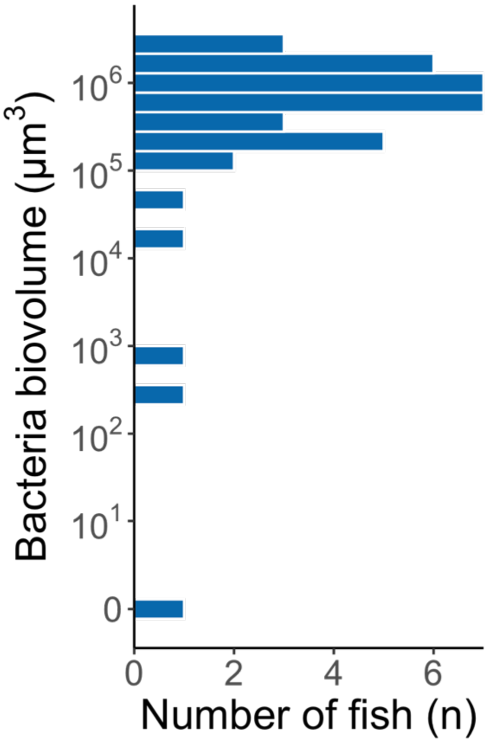


**Figure S1.** Histogram showing distribution of all biovolume measurements (µm$^3$) of large sample of Mix9 gnotobiotic fish (n = 24).

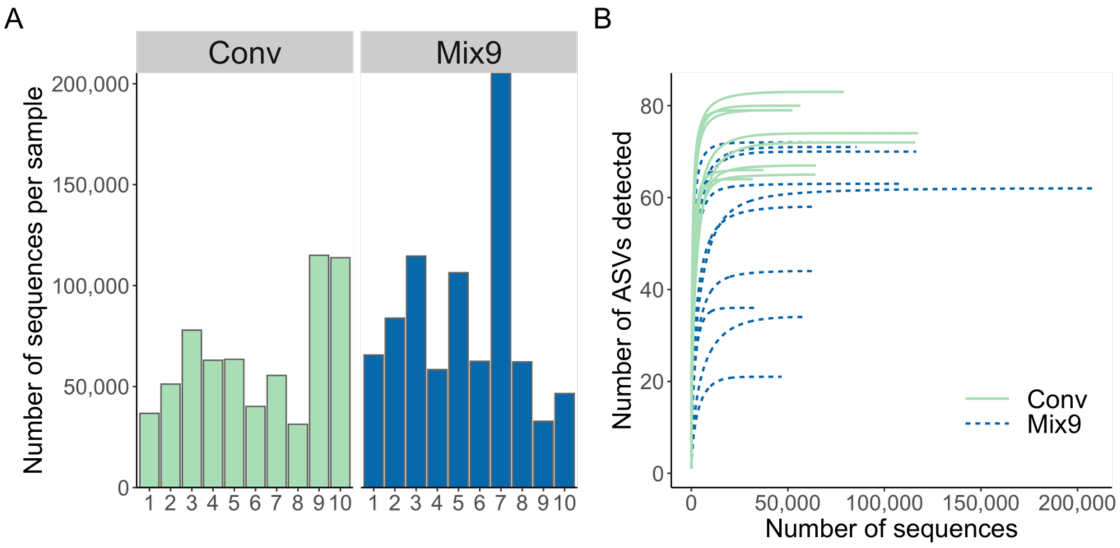


**Figure S2. Quality control metrics for 16S rRNA gene amplicon data. (A)** Number of reads per zebrafish sample. **(B)** ASV rarefaction curves of all zebrafish larvae samples by model type (n = 10 fish per condition).

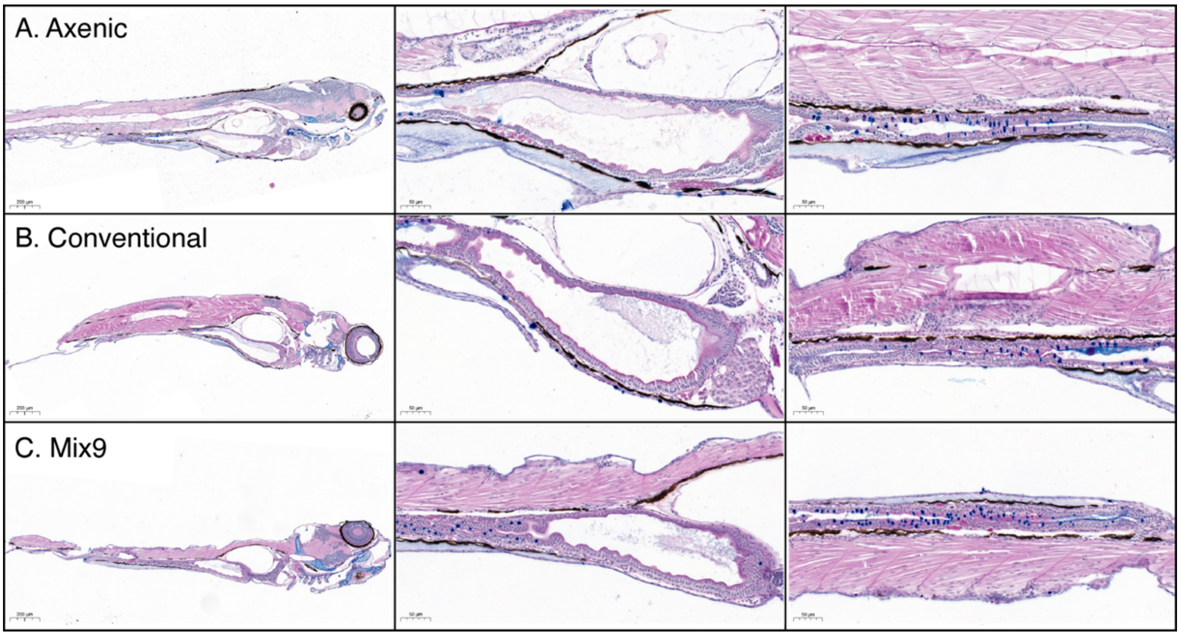


**Figure S3. Histological analysis of larval zebrafish models.** Representative images of the histological analysis of PAS-AB stain of distal intestine of **(A)** Axenic, **(B)** Conventional and **(C)** Mix9 zebrafish at 7 dpf. For each condition, a single fish is shown at 5X (left), then bulb (middle) and distal intestine (right) at 20X. Neutral glycoproteins corresponding to mucus layer are stained in magenta and Goblet cells rich in acidic mucins and glycoproteins are stained in dark blue.
